# Quality of Life, Salivary Cortisol and Atopic Diseases in Young Children

**DOI:** 10.1101/570960

**Authors:** Leif Bjarte Rolfsjord, Håvard Ove Skjerven, Egil Bakkeheim, Teresa Løvold Berents, Kai-Håkon Carlsen, Karin C Lødrup Carlsen

## Abstract

**Background:** Children with atopic disease may have reduced health-related quality of life (QoL) and morning cortisol. The link between QoL, cortisol and atopic disease is unclear.

We aimed to determine if QoL was associated with morning salivary cortisol at two years of age, and if asthma, atopic dermatitis and/or allergic sensitisation influenced this association. Secondarily, we aimed to determine if QoL at one year of age was associated with salivary cortisol one year later.

**Methods and findings:** From the Bronchiolitis All SE-Norway study, enrolling infants during hospitalisation for acute bronchiolitis in infancy (bronchiolitis group) and population based control infants (controls), we included all 358 subjects with available Infant Toddler Quality of Life Questionnaire™ (ITQOL) consisting of 13 domains, and morning salivary cortisol measurements at two years of age. Additionally, QoL nine months after enrolment was available for 289 of these children at one year of age. Recurrent bronchial obstruction was used as an asthma proxy. Atopic dermatitis was defined by Hanifin and Rajka criteria and allergic sensitisation by a positive skin prick test. Associations between QoL and cortisol were analysed by multivariate analyses, stratified by bronchiolitis and control groups due to interaction. At two years of age, QoL was significantly associated with 8/13 QoL domains in the bronchiolitis group, but only with General health in the controls. The associations in the bronchiolitis group showed 0.06-0.19 percentage points changes per nmol/L cortisol for each of the eight domains (p-values 0.0001-0.034). The associations for all domains remained significant, but were diminished by independently including recurrent bronchial obstruction and atopic dermatitis, but remained unchanged by allergic sensitisation.

In the bronchiolitis group only, 8/13 age and gender adjusted QoL domains in one-year old children were significantly associated with cortisol levels at two years (p= 0.0005-0.04).

**Conclusions:** At two years, most QoL domains were associated with salivary cortisol in children who had been hospitalised for acute bronchiolitis in infancy, but for one domain only in controls. The associations were weakened, but remained significant by taking into account asthma and atopic dermatitis. The QoL in one-year old children was associated with salivary cortisol 10 months later.

## INTRODUCTION

Reduced health related quality of life (QoL) has been reported in children with asthma (1, 2) and atopic dermatitis (AD) (3) as well as in infants admitted to hospital for acute bronchiolitis (4-8), a disease known to increase the risk of later asthma (9, 10). Also, the generic Infant Toddler Quality of Life Questionnaire™ (ITQOL) has shown reduced QoL in young children with obstructive airways disease (11), AD (7) and other diseases (5), while reduced QoL also may be associated with psychological and physical stress (12).

Acute bronchiolitis may represent acute stress to the infant, as we recently showed that infants with moderate to severe acute bronchiolitis had higher morning salivary cortisol than healthy infants (13), in line with the higher morning salivary cortisol observed in children and adults during acute stress (14). Furthermore, the severity of acute bronchiolitis has been associated with plasma cortisol suppressing the T-helper cell type 1 (Th1) immune response (15), possibly leading to a shift to a Th2 response in acute bronchiolitis through inhibition of the interferon gamma response acting directly on T cells or indirectly through IL-12. Glucocorticoids may stimulate the secretion of IL-4 and IL-10, enhancing the Th-2 response, and stimulate Th2-cells directly (16), as well as possibly supressing Th2-inflammation (17).

The reduced basal morning cortisol levels observed in children with asthma, also without concurrent use of inhaled corticosteroids (ICS) (18), may on the other hand indicate chronic immunological stress. The subsequent blunted cortisol responses to acute stress in subjects with asthma related to a disturbance of the hypothalamus-pituitary-adrenal (HPA) axis differs from the chronic stress in non-atopic children that can lead to a higher cortisol response (19, 20). Similarly, a blunted cortisol response to acute stress has been suggested in children with AD (21). On the other hand, a higher morning salivary cortisol was observed in two year old children with AD and allergic sensitisation, while a non-significant tendency to lower cortisol was found in children with at least three wheeze episodes (22).

In adults with poorly controlled asthma, both reduced morning salivary cortisol as well as reduced QoL have been found (23), while possible links between QoL, cortisol and atopic disease in children are not known.

Morning salivary cortisol, reflecting the non-protein bound fraction of serum cortisol (24, 25) was recently shown to be similar in two year old children with and without acute bronchiolitis in infancy (13) with values ranging from 2.5 to 189.0 nmol/L, and with higher levels in girls than in boys.

Based upon our previous findings of high morning cortisol levels during acute bronchiolitis but with levels at two years of age similar to those of controls (13), we hypothesised that *low* cortisol levels in periods without acute illness may contribute to development of asthma. We further hypothesised that reduced QoL some months after severe acute illness in early life may be a marker of chronic stress, with subsequent lower future salivary cortisol levels.

We therefore primarily aimed to determine if QoL was associated with morning salivary cortisol at two years of age, and if asthma, atopic dermatitis and/or allergic sensitisation modified this association. Secondarily, we aimed to determine if QoL at one year of age was associated with salivary cortisol at two years.

## MATERIALS AND METHODS

### Study design

From the source population of 644 children included in the Bronchiolitis ALL SE-Norway study enrolling infants who were hospitalised for acute bronchiolitis and controls recruited from a general population (7), we included all 358 children with available salivary cortisol and QoL at 24 months of age. The bronchiolitis group consisted of 203 infants with moderate to severe acute bronchiolitis at inclusion, and 155 were controls. For details, see figure 1 and Supporting Information.

**Legend Figure 1:**
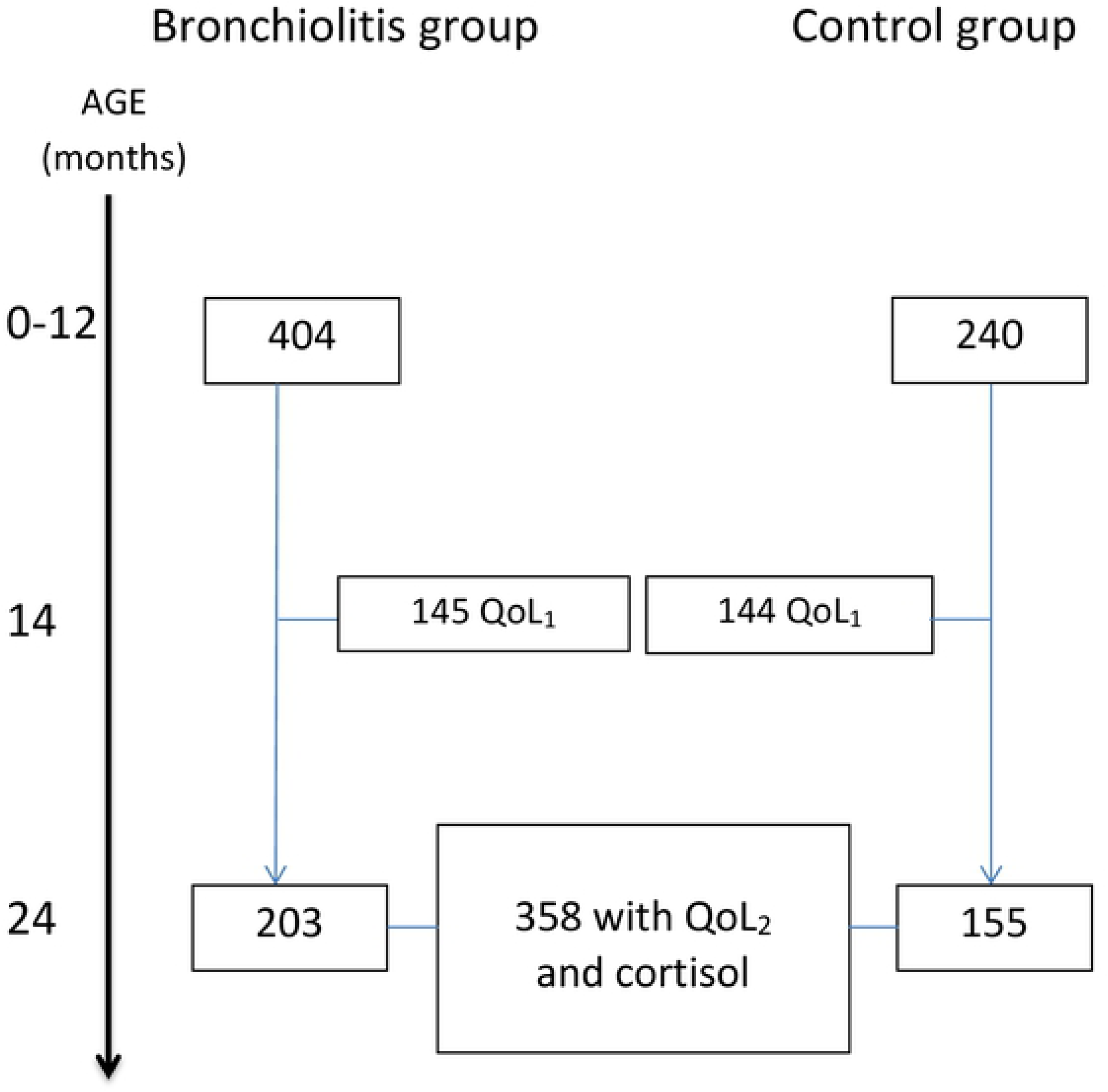
The figure outlines the number of infants enrolled in the Bronchiolitis All SE-Norway study (top, n=644) who were subsequently included in the present study (n=358) for analyses based upon available Quality of life (QoL) and/or salivary morning cortisol at 24 months of age. The QoL questionnaires were completed nine months after enrolment at approximately 14 months of age (QoL_1_) as well as at the time of the clinical examination at 24 months of age (QoL_2_).

Investigations at enrolment and at two years of age included clinical assessment, structured parental interviews and morning salivary sampling for cortisol, whereas skin prick test (SPT) for common inhalant and food allergens was performed at two years only. Quality of life questionnaires were completed nine months after enrolment (QoL_1_) (7, 8) and at two years of age (QoL_2_).

Caregivers of all children signed the informed written consents prior to study enrolment. The study was approved by the Regional Committee for Medical and Health Research Ethics and The Norwegian Data Protection Authority and registered in the Norwegian bio bank registry. The randomised clinical trial part of the study was registered in Clinical Trials.gov, no. NCT00817466 (26).

### Study subjects

The mean (range) age of the 358 children in the present study was 5.2 (0.2-13.4) months at enrolment and 24.2 (17.6-34.7) months at the two-year investigation. The children in the bronchiolitis group compared to controls where shorter, more often exposed to second-hand smoke at inclusion and their parents had lower income, lower educational attainment and less often allergic rhinitis or AD (Table 1).

**Table 1.**
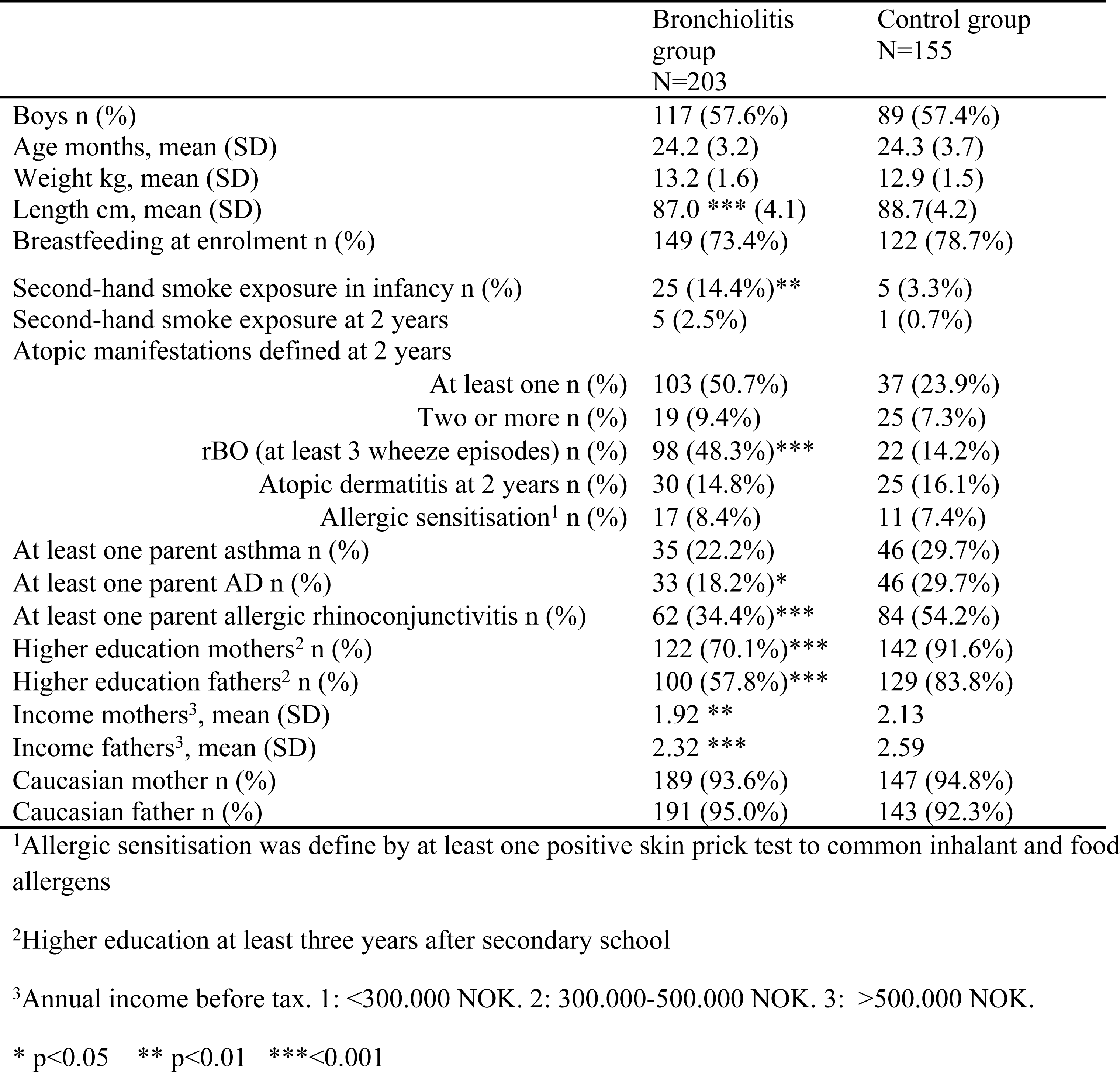
Characteristics and asthma risk factors of the children at two years of age. All data are given as n and %, unless otherwise stated. The control group is the group from the reference population.

|  | Bronchiolitis group<br>N=203 | Control group<br>N=155 |
| --- | --- | --- |
| Boys n (%) | 117 (57.6%) | 89 (57.4%) |
| Age months, mean (SD) | 24.2 (3.2) | 24.3 (3.7) |
| Weight kg, mean (SD) | 13.2 (1.6) | 12.9 (1.5) |
| Length cm, mean (SD) | 87.0 *** (4.1) | 88.7(4.2) |
| Breastfeeding at enrolment n (%) | 149 (73.4%) | 122 (78.7%) |
| Second-hand smoke exposure in infancy n (%) | 25 (14.4%)** | 5 (3.3%) |
| Second-hand smoke exposure at 2 years | 5 (2.5%) | 1 (0.7%) |
| Atopic manifestations defined at 2 years |  |  |
| At least one n (%) | 103 (50.7%) | 37 (23.9%) |
| Two or more n (%) | 19 (9.4%) | 25 (7.3%) |
| rBO (at least 3 wheeze episodes) n (%) | 98 (48.3%***) | 22 (14.2%) |
| Atopic dermatitis at 2 years n (%) | 30 (14.8%) | 25 (16.1%) |
| Allergic sensitisation<sup>1</sup> n (%) | 17 (8.4%) | 11 (7.4%) |
| At least one parent asthma n (%) | 35 (22.2%) | 46 (29.7%) |
| At least one parent AD n (%) | 33 (18.2%*) | 46 (29.7%) |
| At least one parent allergic rhinoconjunctivitis n (%) | 62 (34.4%***) | 84 (54.2%) |
| Higher education mothers<sup>2</sup> n (%) | 122 (70.1%***) | 142 (91.6%) |
| Higher education fathers<sup>2</sup> n (%) | 100 (57.8%***) | 129 (83.8%) |
| Income mothers<sup>3</sup>, mean (SD) | 1.92 ** | 2.13 |
| Income fathers<sup>3</sup>, mean (SD) | 2.32 *** | 2.59 |
| Caucasian mother n (%) | 189 (93.6%) | 147 (94.8%) |
| Caucasian father n (%) | 191 (95.0%) | 143 (92.3%) |

<sup>1</sup>Allergic sensitisation was define by at least one positive skin prick test to common inhalant and food allergens
<sup>2</sup>Higher education at least three years after secondary school
<sup>3</sup>Annual income before tax. 1: <300.000 NOK. 2: 300.000-500.000 NOK. 3: >500.000 NOK.
\* p<0.05 \*\* p<0.01 \*\*\*<0.001

## Methods

Atopic manifestations determined at two years of age, consisted of recurrent bronchial obstruction (rBO) as a proxy for asthma, atopic dermatitis and allergic sensitisation.

*Recurrent bronchial obstruction* was defined as at least three parentally reported episodes of wheeze at two years of age, in line with previous reports (27), with acute bronchiolitis included as one episode. *Atopic dermatitis* was defined based upon the modified Hanifin and Rajka’s criteria (yes or no) (28), and severity by the SCORing AD index (SCORAD) (see Supporting Information for details).

*Allergic sensitisation*, determined by SPT using 17 common inhalant and food allergens with Soluprick SQ allergen extracts, ALK, Hørsholm, Denmark, was defined as positive with at least one mean wheal diameter at least 3 mm greater than the negative control. Further details are given in the Supporting Information.

*Morning salivary cortisol* was sampled by the parents on the first morning after enrolment in the bronchiolitis group, otherwise at home and brought to the investigation centre. Two Sorbette® hydrocellulose microsponges were applied in the child’s mouth as soon as possible after their child’s first awakening after 6:00 a.m., before the first meal, and placed in appropriate prepared containers, as described elsewhere (13). The samples were stored at −86°C and later analysed at Karolinska Institutet, Stockholm, with radioimmunoassay with monoclonal rabbit antibodies Codolet, France).

*The Infant Toddler Quality of Life™ Questionnaire* (*IT*_*QOL*_*-97*) (11) completed by the parents included 97 questions within 13 domains scored from 0 (worst) to 100 (best), with no overall score. Accordingly, a change in QoL score is equivalent to the percentage point score change. The Overall health domain consisted of only one item: Is your child’s health excellent, very good, good, fair or poor? In line with others (29) and as previously reported (7, 8), with permission from the copyright holder, we recoded the domain Change in health from the original scores from 1-5 to 0-100 (zero meaning worst deterioration of health from one year ago, 50 meaning no change). Four domains (Change in health, General behaviour, Overall behaviour and Getting along) were recorded in children older than 12 months only (7).

### Study outcomes and explanatory variables

The main outcome for our primary aim, QoL_2_, was reported by quantitative values per domain, and secondarily by the number of domains with significantly reduced QoL_2_ scores.

The main explanatory variables for the primary aim were morning salivary cortisol, and the three atopic manifestations rBO, AD and allergic sensitisation at two years of age. Further analyses reported in Supporting Information substituted the respective atopic manifestations by quantitative measures, i.e. the total number of wheeze episodes, the AD severity score SCORAD and the sum of SPT wheal diameters for influence on the associations between morning salivary cortisol and QoL_2_.

The main outcome of the secondary aim was morning salivary cortisol, with QoL_1_ as the explanatory factor.

### Statistical analysis

The bronchiolitis and control groups were compared by Pearson’s chi-square tests for categorical data and Student’s T-test for normally distributed numerical data, and otherwise with Welch test, unless otherwise stated.

Due to non-normality of results and residuals, we used linear robust regression by Huber’s M method (30), for analyses including QoL and cortisol. To estimate the relative influence by rBO, AD and allergic sensitisation on QoL_2_, we calculated the percent point change equivalent to the difference in score for each QoL domain, given per nmol/L change in cortisol. For comparison, we calculated the difference in each QoL domain score that was attributed to a difference in salivary cortisol level of 95th versus 5^th^percentile (QoL score at the salivary cortisol level of 95^th^percentile minus QoL score at the 5^th^percentile). Salivary cortisol was studied as a continuous variable, and presented graphically by quartiles.

Each atopic manifestation was included in robust regression models to assess their potential influence on both cortisol and QoL_2_, as well as the associations between the two (see Figure 2, hypothesis). For graphical presentations of QoL versus cortisol levels and cortisol levels versus atopic manifestations we used data unadjusted for age and gender. In line with previously demonstrated associations between morning salivary cortisol and age as well as gender (13), we decided a priori to analyse age and gender adjusted associations between cortisol and QoL as well between QoL and atopic manifestations. The atopic manifestations were not considered to be confounders, as they could be causally associated with both cortisol and QoL_2_.

**Legend Figure 2:**
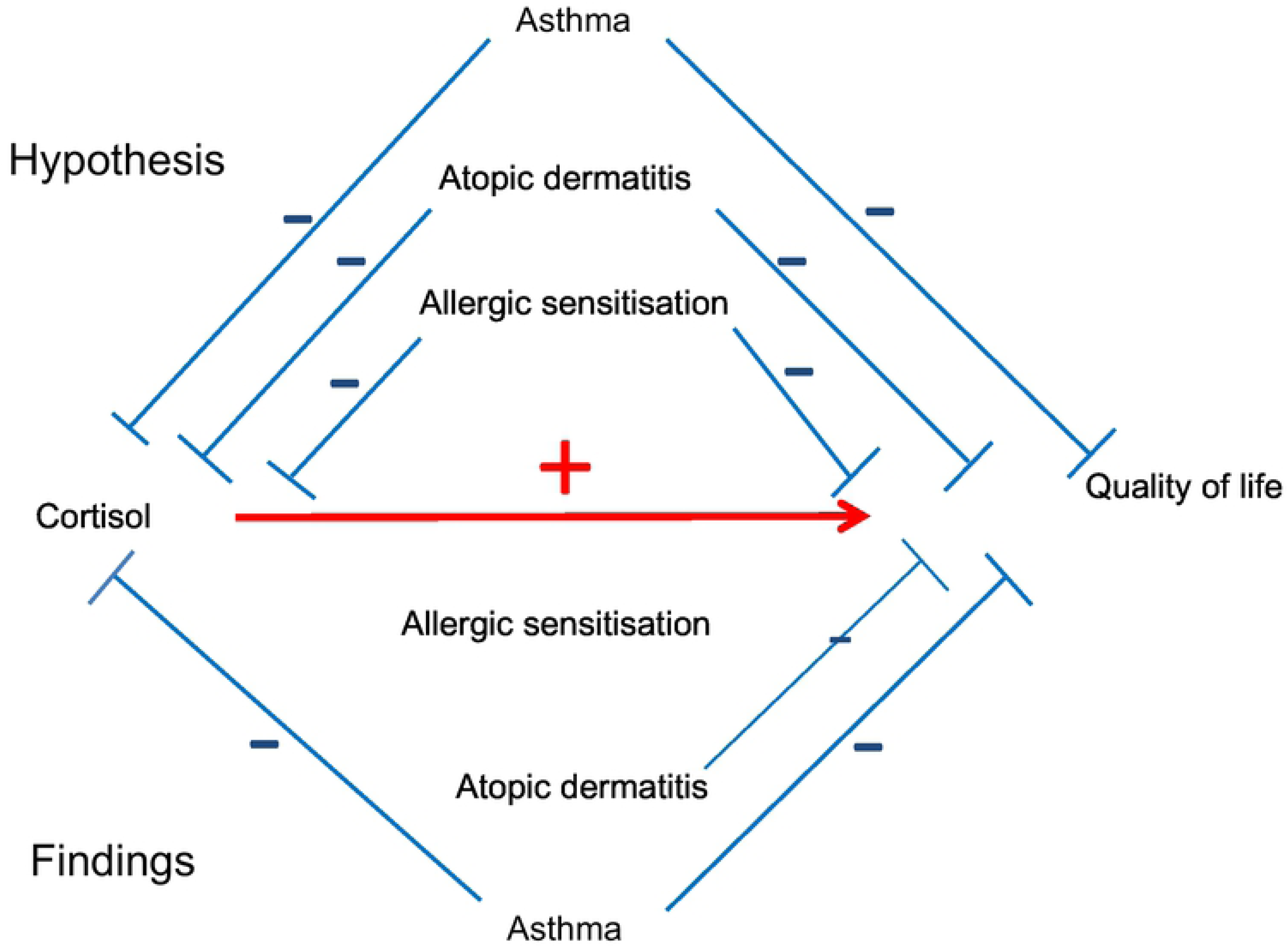
Directed acyclic graph showing hypothesised influence on cortisol and QoL24m by allergic diseases above the red line, and observed influence in the bronchiolitis group below the red line. The red line indicates the net result from the influence of allergic disease on the association between morning salivary cortisol and QoL24m

Using QoL_2_ as dependent variable in two-way regression analyses, we tested for interactions between the group affiliation (bronchiolitis or controls) and cortisol, as well as between atopic manifestations and cortisol. Due to interactions between group affiliation and salivary cortisol as well as atopic dermatitis, analyses were stratified by group affiliation.

Possible confounding was assessed by robust regression and considered relevant if the outcome of the model was changed by at least 25% (31) by any of the possible confounders (socioeconomic factors, parental allergic disease, secondary smoke). Confounding by socioeconomic factors was tested by including these factors in multiple regression models, and eliminating the factors with highest p-values stepwise by Hosmer’s procedure (31) until only factors with p-values < 0.05 remained, retaining age and gender.

The level of statistical significance was set to p<0.05 for all analyses.

Analyses were performed with the IBM SPSS Statistics 21 (IBM Corporation, Armonk, New York, USA), and the NumberCruncher Statistical System (NCSS Kaysville, Utah, USA), version 11.

## RESULTS

### Atopic manifestations and QoL_2_; bronchiolitis group vs. controls

Children in the bronchiolitis group were significantly more often affected by at least one atopic manifestation at two years and had more often rBO than the controls, while AD was similar in the two groups (Table 1).

The QoL_2_ scores varied from 0-100 in five domains, with the smallest score range seen in the domain Getting along (53.3), as shown in Table 2. The bronchiolitis group reported larger improvement in health (Change in health), while controls scored significantly higher for Overall health and General health (Table 2).

**Table 2.** Unadjusted weighted means (95% CI) of QoL at two years of age (QoL24m) of children included at hospitalisation for acute bronchiolitis and control children

| Domain | Previous bronchiolitis | Control children | All children |
| --- | --- | --- | --- |
|  | Unadjusted weighted means (95% CI) |  | Median (min, max) |
| Overall health | 83.4 (81.3, 85.5)** | 88.7 (86.3, 91.1) | 85.0 (0.0, 100.0) |
| Physical abilities | 100.0 (100.0, 100.0) | 100.0 (100.0, 100.0) | 100.0 (0.0, 100.0) |
| Growth and development | 94.7 (93.7, 95.6) | 95.2 (94.1, 96.3) | 97.2 (0.0, 100.0) |
| Bodily pain/ discomfort | 80.5 (78.4, 82.6) | 78.5 (76.0, 80.9) | 75.0 (8.3, 100.0) |
| Temperament and moods | 84.2 (83.0, 85.4) | 83.0 (81.7, 84.4) | 84.7 (36.8, 100.0) |
| General behaviour | 84.5 (82.9, 86.0) | 85.2 (83.4, 86.9) | 85.4 (35.4, 100.0) |
| Overall behaviour | 85.0 (85.0, 85.0) | 85.0 (85.0, 85.0) | 85.0 (30.0, 100.0) |
| Getting along | 78.8 (77.6, 80.0) | 78.5 (77.2, 79.9) | 78.3 (45.0, 98.2) |
| General health | 67.1 (65.0, 69.2)**** | 78.3 (75.9, 80.7) | 75.0 (18.2, 100.0) |
| Change in health | 65.2 (62.8, 67.8)**** | 56.9 (54.1, 59.7) | 50.0 (0.0, 100.0) |
| Parental impact – emotions | 91.3 (90.1, 92.6) | 91.3 (89.9, 92.7) | 92.9 (35.7, 100.0) |
| Parental impact – time | 95.2 (94.2, 96.1) | 94.2 (93.1, 95.3) | 95.2 (28.6, 100.0) |
| Family cohesion | 79.6 (77.3, 82.0) | 80.8 (78.1, 83.5) | 85.0 (0.0, 100.0) |

\*\*\*\*p<0.0001

### Quality of life and salivary cortisol at two years of age

In the bronchiolitis group, eight QoL_2_ domains were significantly and positively associated with morning salivary cortisol (p=0.0001-p=0.035), see Table 3 and Figure 3. The association between Overall health and salivary cortisol was significant only in boys, (p<0.0001).

**Legend Figure 3:**
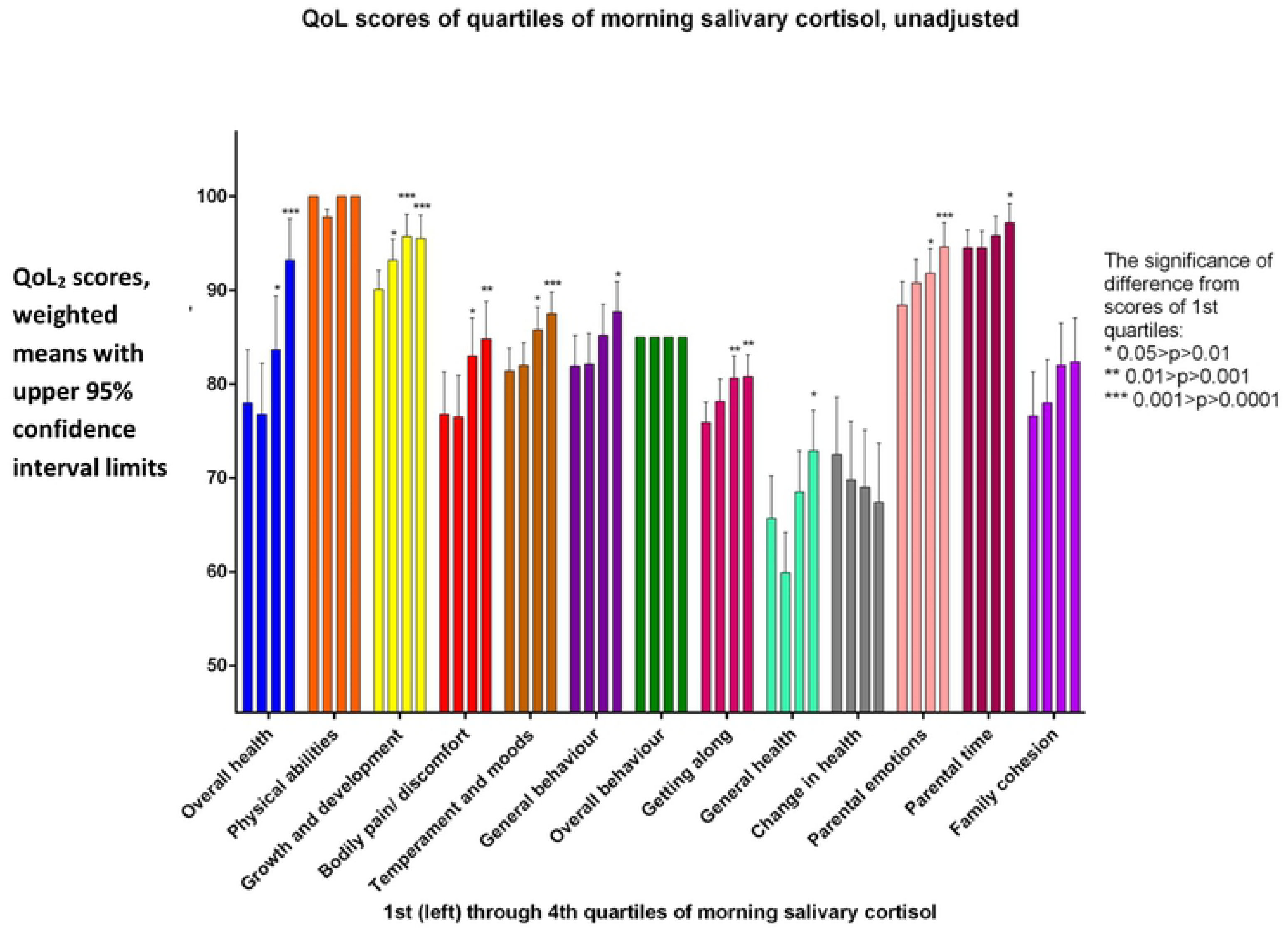
Bronchiolitis group: QoL_2_ scores for each domain, unadjusted, for each quartile of morning salivary cortisol, 1^st^quartile lowest cortisol, 4^th^quartile highest. Due to interaction between gender and cortisol for the Overall health domain, this domain was analysed separately for the genders. An association was found only for boys for this domain. For Overall health, results for boys are shown. For the other domains, results for both genders analysed together are shown.

**Table 3.** The potential influence of recurrent bronchial obstruction (rBO), atopic dermatitis (AD) and allergic sensitisation (AS) on the associations between Quality of Life (QoL_2_) and salivary cortisol at two years of age is shown for 203 children who had moderate to severe acute bronchiolitis in infancy. The influence by including each atopic manifestation (rBO, AD and AS) is shown as the percentage change of QoL per 1 nmol/L change in salivary cortisol, adjusted for age and gender. Each column includes all children with the observed atopic manifestation, and they are not mutually exclusive.

| Domain (Mean domain score difference<sup>1</sup><br>by difference between 95<sup>th</sup> and 5<sup>th</sup><br>percentile of cortisol, 51.6 nmol/L) | Change in QoL<sub>2</sub><br>score per nmol/L<br>unit salivary cortisol | rBO<br>% change in association | AD | Allergic<br>sensitisation |
| --- | --- | --- | --- | --- |
| Overall health<sup>2</sup> boys (16.0) | 0.31 (0.17, 0.45)**** | -20.7 | -2.1 | -0.5 |
| Overall health girls (-0.0) | -0.00 (-0.16, 0.16) |  |  |  |
| Growth and development (3.8) | 0.07 (0.02, 0.13)** | -1.4 | 1.7 | 0.6 |
| Bodily pain/ discomfort (6.2) | 0.12 (0.01, 0.23)* | -8.3 | 5.4 | -2.3 |
| Temperament and moods (6.1) | 0.12 (0.06, 0.18)*** | -6.9 | 1.4 | -1.5 |
| General behaviour (4.6) | 0.09 (0.01, 0.17)* | -6.7 | -1.5 | 1.3 |
| Getting along (4.0) | 0.08 (0.02, 0.13)** | -3.0 | -0.3 | 0.9 |
| General health (5.6) | 0.11 (-0.00, 0.22)<sup>3</sup> | -26.9 | 0.4 | 0.3 |
| Parental impact –Emotions (4.5) | 0.09 (0.3, 0.15)** | -6.1 | -5.1 | 1.2 |
| Parental impact – Time (3.0) | 0.06 (0.01, 0.10)* | -7.7 | -0.1 | 0.9 |

<sup>1</sup>QoL score difference equals percentage point difference.
<sup>2</sup>Overall health was gender stratified due to interaction.
\* $p<0.05$ \*\* $p<0.01$ \*\*\* $p<0.001$ \*\*\*\* $p<0.0001$
<sup>3</sup> $p=0.0517$

In the controls, General health only was significantly associated with cortisol. The significant increase of 0.1 percentage point per nmol/L in cortisol level (95% CI 0.0, 0.2, p=0.046) corresponded to a QoL_2_ difference of five percent points between children having cortisol levels at the 5^th^vs 95^th^percentile (a difference of 51.6 nmol/L of salivary cortisol). No further analyses were performed in this group, with only one QoL domain significantly associated with salivary cortisol.

The hypothesised (top) and observed (bottom) influence of atopic manifestations on cortisol and QoL_2_ are shown schematically in Figure 2. As shown in Table 2, the strongest influence on the associations between cortisol and QoL_2_ was exerted by rBO, reducing the associations with 1.4 to 26.9 per cent, followed by changes related to AD ranging from −5.5 to 5.1 and less than 3 per cent changes by allergic sensitisation. However, all associations between QoL and cortisol remained significant after including rBO, AD and allergic sensitisation into the regression analyses.

Finally, we found no significant confounding effect of socioeconomic factors, parental ethnicity and second-hand smoke at two years of age, and these were consequently not included in the final multivariate analyses (see Supporting Information, Table 4 for details).

**Table 4.**
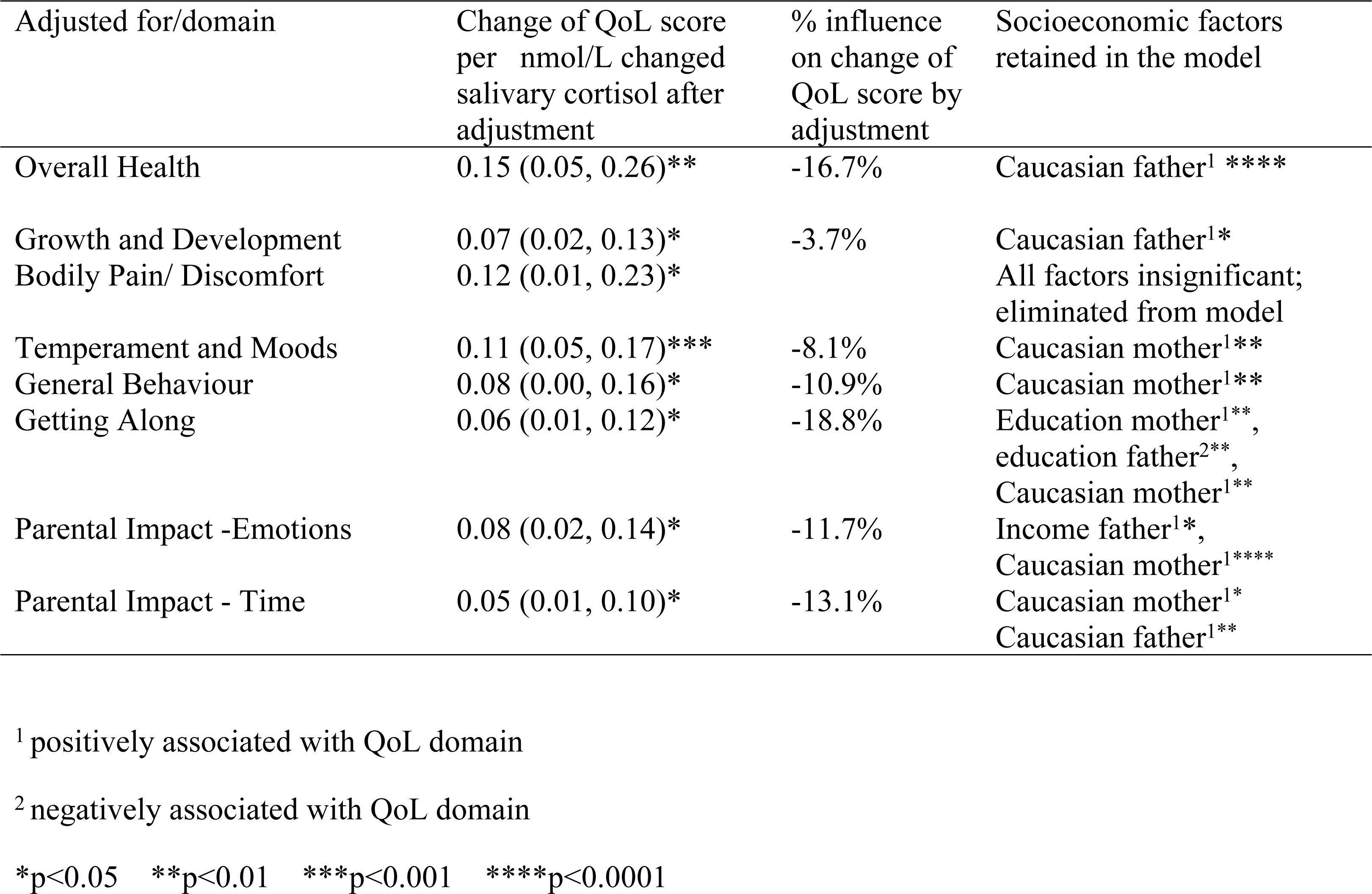
Bronchiolitis group: Change of associations between salivary morning cortisol at and QoL24m at two years of age by socioeconomic factors, including age, gender, and the following socioeconomic factors: mother’s education, father’s education, mother’s income, father’s income, ethnicity of father and of mother (Caucasian or not) and secondhand smoke exposure at two years of age. The socioeconomic factors have been eliminated by Hosmer’s stepdown procedure, finally retaining factors with p<0.05. Age and gender have been retained in the models

### Salivary cortisol in bronchiolitis group vs. controls, and atopic manifestations

The age and gender adjusted salivary cortisol levels at two years were similar in the bronchiolitis group and controls. Weighted mean difference was −0.70 (95% CI −3.7, 2.3) nmol/L.

Salivary cortisol was significantly lower in children with rBO vs. children without rBO for the bronchiolitis group and controls together (weighted mean difference −4.1 (95%CI −7.3,-1.0) nmol/L), as shown schematically in Figure 2, and in unadjusted analysis in Figure 4. Neither AD nor allergic sensitisation was significantly associated with morning salivary cortisol at two years of age (Figures 1 and 3).

**Legend Figure 4:**
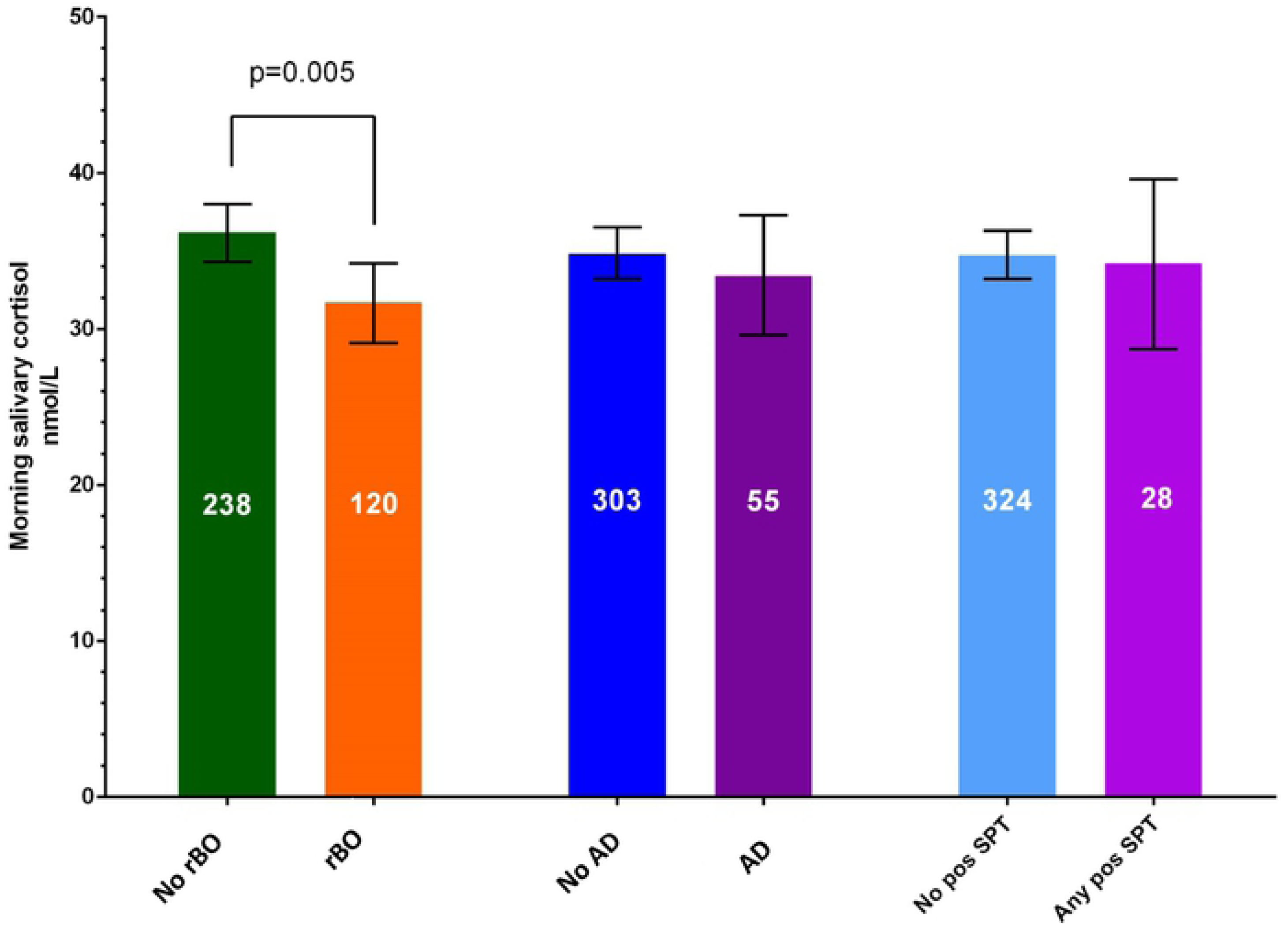
Morning salivary cortisol in children of the bronchiolitis group and controls together, without recurrent bronchial obstruction (rBO), defined as at least three wheeze episodes, compared to with rBO, no atopic dermatitis (AD) vs. with AD as well as with no or any positive skin prick test (SPT) to common inhalant and food allergens.

### QoL_2_ and atopic manifestations

The QoL_2_ was significantly associated with rBO and AD in the bronchiolitis group, and with rBO as well as allergic sensitisation in the controls, as shown in Table 5, referring to absolute changes (= percentage point changes) in QoL_2_ scores by the atopic diseases.

**Table 5.** The impact of allergic diseases on QoL_2_ is given for each domain as mean (95% CI), adjusted for age and gender, given for infants with moderate to severe bronchiolitis in infancy compared to controls. As an example; the negative association of General health (GH) with rBO is stronger in the bronchiolitis group than among controls, both being statistically significant.

|  | Recurrent bronchial obstruction |  | Atopic dermatitis |  | Allergic sensitisation |  |
| --- | --- | --- | --- | --- | --- | --- |
|  | bronchiolitis group | controls | bronchiolitis group | controls | bronchiolitis group | controls |
| OH | -12.3 (-16.5, -8.2)**** | -6.6 (-13.9, 0.7) | -5.5 (-11.7, 0.6) | -6.3 (-13.6, 0.9) | -0.7 (-8.6, 7.2) | -11.7 (-21.6, -1.9)* |
| PA | -1.2 (-2.1, -0.4)** | Not done<sup>3</sup> | -0.0 (-0.0, 0.0) | -2.1 (-3.2, -1.0)*** | 0.0 (-0.2, 0.2) | Not done<sup>7</sup> |
| GD | -3.5 (-5.8, -1.2)** | -1.3 (-4.3, 1.8) | -2.5 (-5.8, 0.7) | -1.2 (-4.2, 1.8) | -0.1 (-4.3, 4.0) | -1.9 (-6.4, 2.6) |
| BD | -6.5 (-10.8, -2.1)** | -7.3 (-14.7, 0.1) | -7.2 (-13.4, -0.9)* | -1.7 (-8.6, 5.3) | 4.2 (-3.6, 12.1) | -6.4 (-16.8, 4.0) |
| TM | -3.1 (-5.5, -0.7)* | -4.1 (-7.8, -0.5)* | -5.1 (-8.4, -1.8)** | 1.0 (-2.6, 4.5) | -2.5 (-1.8, 6.8) | -1.3 (-6.5, 3.9) |
| GB | -4.7 (-7.9, -1.4)** | -0.7 (-5.5, 4.1) | -5.3 (-9.9, -0.7)* | -4.1 (-8.8, 0.6) | -0.8 (-6.8, 5.2) | -7.0 (-13.7, -0.3)* |
| OB | -0.0 (-0.0, 0.0) | Not done<sup>4</sup> | 0.0 (0.0, 0.0) | Not done<sup>5</sup> | 0.0 (-0.0, 0.0) | Not done<sup>7</sup> |
| GA | -1.9 (-4.2, 0.4) | -4.0 (-7.9, -0.0)* | -6.2 (-9.3, -3.0)*** | -0.9 (-4.8, 2.9) | -0.4 (-4.6, 3.8) | -8.0 (-13.6, -2.4)** |
| GH | -13.8 (-17.8, -9.8)**** | -6.8 (-12.9, -0.7)* | -5.8 (-12.0, 0.3) | -7.0 (-13.0, -1.0)* | 0.1 (-7.9, 8.1) | -12.2 (-20.5, -3.9)** |
| CH | 6.9 (0.8, 13.0)* | 6.8 (-0.6, 14.2) | 7.8 (-0.9, 16.4) | 3.1 (-3.3, 9.6) | -1.0 (-12.3, 10.3) | 7.3 (-2.8, 17.4) |
| PE | -4.0 (-6.4, -1.5)** | -2.3 (-5.9, 1.4) | -5.7 (-9.2, -2.2)** | -1.6 (-5.2, 2.0) | -0.2 (-4.7, 4.3) | -5.8 (-10.7, -1.0)* |
| PT | -2.5 (-4.5, -0.5)* | -2.1 (-5.0, 1.0) | -1.9 (-4.8, 1.0) | -1.9 (-4.9, 1.0) | -0.0 (-3.7, 3.7) | -4.3 (-8.3, -0.2)* |
| FC | -0.1 (-4.7, 4.5) | 0.3 (-7.4, 8.0) | -5.4 (-11.9, 1.1) | 0.7 (-6.7, 8.1) | -0.0 (-3.7, 3.7) | -1.8 (-12.3, 8.7) |

<sup>3</sup>Not done because gender is highly correlated with other X's <sup>4</sup>Not done because rBO is highly correlated with other X's <sup>5</sup>Not done because AD is highly correlated with other X's <sup>6</sup>Not done, procedure call invalid <sup>7</sup>Not done because AS is highly correlated with other X's
OH = Overall health; PA = Physical abilities; GD = Growth and development; BD = Bodily pain/ discomfort; TM = Temperament and moods; GB = General behaviour; OB = Overall behaviour; GA = Getting along; GH = General health; CH = Change in health; PE = Parental impact – emotions; PT = Parental impact – time; FC = Family cohesion

### Association between QoL_1_ and salivary cortisol at two years

In the bronchiolitis group only, QoL_1_ was significantly and positively associated with morning salivary cortisol at two years of age in age and gender adjusted analysis for 8/13 domains, as shown in Table 6.

**Table 6.**
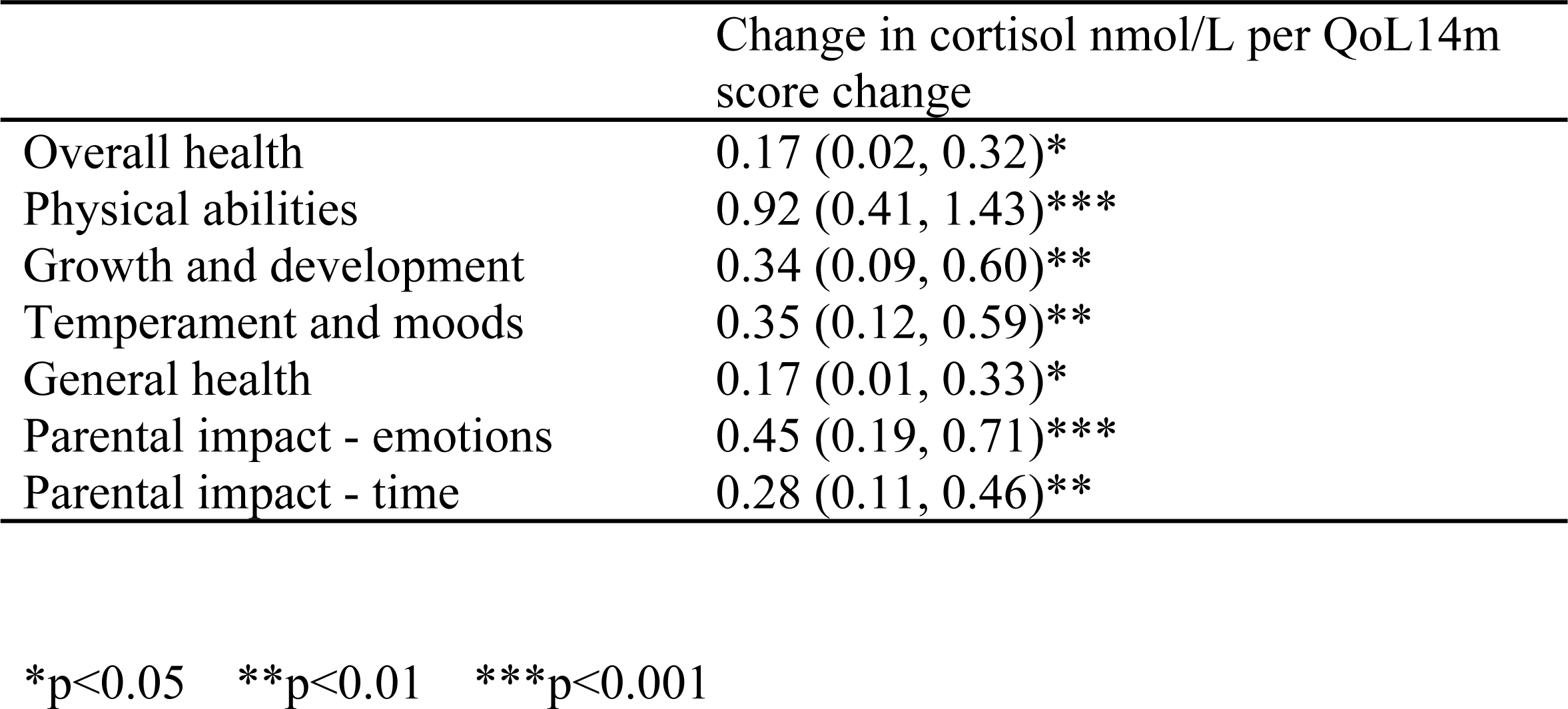
Associations between QoL_1_ (quality of life at a mean age of 14 months) scores and subsequent cortisol at visit 2, at a mean age of 24 months, bronchiolitis group only, adjusted for age at the two year old visit and gender.

|  | Change in cortisol nmol/L per QoL14m score change |
| --- | --- |
| Overall health | 0.17 (0.02, 0.32)* |
| Physical abilities | 0.92 (0.41, 1.43)*** |
| Growth and development | 0.34 (0.09, 0.60)** |
| Temperament and moods | 0.35 (0.12, 0.59)** |
| General health | 0.17 (0.01, 0.33)* |
| Parental impact - emotions | 0.45 (0.19, 0.71)*** |
| Parental impact - time | 0.28 (0.11, 0.46)** |

\*p<0.05 \*\*p<0.01 \*\*\*p<0.001

## DISCUSSION

Quality of life was significantly associated with morning salivary cortisol at two years of age for most domains among children hospitalised with acute bronchiolitis in infancy, i.e. low cortisol was associated with low QoL, and high cortisol with high QoL. Significant associations were observed for the General Health domain only in controls. The associations remained significant, but were weakened by rBO, with only limited or no significant influence by AD or allergic sensitisation. The QoL in one-year old children was associated with salivary cortisol at two years of age.

We are not aware of other studies comparing QoL and morning salivary cortisol in children. Our findings that after moderate to severe acute bronchiolitis in infancy, QoL_2_ was associated with salivary cortisol at two years of age, is to our knowledge novel. The results could not be confirmed in the population based controls. We have previously found that infants with acute bronchiolitis have higher morning salivary cortisol than controls (13), indicating acute stress. Others have found other signs of acute stress in acute bronchiolitis with respiratory syncytial virus, differing from other infections and acute diseases (32). Reduced QoL, found after acute bronchiolitis (5, 33), may partly be expressions of chronic stress, not only physical, but possibly psychological stress. Concerns of the parents of the children of the bronchiolitis group, as indicated by the Parental impact – emotions and Parental impact – time domains in the present study, seem to be associated with the children’s cortisol levels. The associations that we found after acute bronchiolitis between cortisol and QoL in domains reflecting expressions of pain, moods and behaviour, i.e. Bodily pain/ discomfort, Temperament and moods, General behaviour and Getting along, partly influenced by rBO and AD, may also indicate a role of psychological stress in the development of atopic disease. Other studies have shown that pre-and postnatal maternal distress and negative life events are associated with allergic disease in children (19, 34). This may suggest long term effects on QoL of severe lower respiratory infection in infancy, possibly through chronic stress which has been negatively associated with morning cortisol (18, 35).

The 16 percentage point difference in Overall health in boys with low versus high salivary cortisol is likely to be clinically relevant as they are comparable to the eight percentage point General health differences between children with and without asthma-like symptoms reported from the Generation R study (36).

The observed association between Overall health and morning salivary cortisol at two years of age was significant among both genders analysed together, but only in boys by gender stratified analyses. This may be explained by our significantly higher salivary cortisol levels in girls compared to boys in the Bronchiolitis ALL study reported previously (13).

The influence by rBO, and to a lesser extent AD, on the associations between QoL and salivary cortisol in children who were hospitalised for acute bronchiolitis in infancy in our study point to complex associations between allergic diseases, QoL and salivary cortisol. This is in line with attenuated regulation of the HPA axis previously shown in children with chronic atopic diseases, with a reduced cortisol after acute stress provocation, indicating chronic stress (16, 19, 35, 37) and reduced morning salivary cortisol observed in adults (23) and children (18) with poor asthma control. Our previous findings of increased morning salivary cortisol during acute bronchiolitis in infancy (13) and reduced QoL nine months later (7), were extended in the present study, showing that rBO, but not one episode of bronchiolitis (13) was associated with lower morning salivary cortisol at two years of age. This may indicate that repeated episodes of bronchial obstruction are necessary to affect cortisol levels, possibly through chronic stress (13, 15, 19), irrespective of current use of inhaled corticosteroids (23). On the other hand, our study was not designed to identify the reverse possibility of a potential causal role of low salivary cortisol levels outside acute respiratory disease for the development of recurrent bronchial obstruction.

The influence of AD on the associations between QoL and cortisol in our study was less clear, possibly because most children with AD had mild disease. However, supporting our findings of some influence on the cortisol-QoL association, Stenius et al. reported higher morning salivary cortisol in two year old children with atopic dermatitis as opposed to children with recurrent wheeze who had a non-significant tendency to lower salivary cortisol (22). The possible opposite association between rBO and AD with cortisol in early childhood is not clear, since both diseases have been associated with reduced QoL. The lack of significant associations between allergic sensitisation and QoL in the bronchiolitis group and allergic sensitisation and salivary cortisol may have several explanations. In our study less than 10 per cent of the subjects were sensitised to at least one allergen, limiting the likelihood of observing significant associations. On the other hand, allergic sensitisation may not affect QoL before allergen exposure causes symptoms, which for inhalant allergens occur more frequently with increasing age (38).

Our finding that reduced QoL about one year of age was associated with lower salivary cortisol at two years of age is to our knowledge also novel. We recently showed in the same study population that in addition to having been hospitalised for acute bronchiolitis, disease severity and asthma risk factors as well as AD were associated with reduced QoL at 14 months of age (7, 8).

The direct clinical implications of our findings remain unclear at present. The influence by our asthma proxy of rBO dominated the association between QoL and salivary cortisol, weakening most of the associations by more than five per cent. This finding is supported by a stronger influence by the total number of wheeze episodes, as shown in the Supporting Information. The maintenance of statistical significance of the influence of cortisol on QoL after including rBO in the regression model also indicates an additive negative effect of low cortisol and rBO on QoL. The implications in terms of the Overall health domain could be shown by a 24-months-old boy with rBO and low salivary cortisol, at the 5^th^percentile, having an estimated 23.1 percentage point lower QoL than a boy without rBO who had a high salivary cortisol level, at the 95^th^percentile. However, our study suggests that in addition to rBO and to some extent AD, also acute moderate to severe infant bronchiolitis may play a role in the association between future salivary cortisol and QoL. Although acute infant bronchiolitis per se may not lead to future low cortisol levels, QoL was reduced both at 14 and 24 months of age compared to controls. Although the influence of the asthma proxy of at least three episodes of bronchial obstruction in our study dominated the association between cortisol and QoL at two years, the associations were significant also among children in the bronchiolitis group who did not go on to develop rBO, as shown in Supporting Information. Together these observations suggest that children who have acute bronchiolitis in infancy and reduced QoL some months later may be at increased risk of chronic stress, observed by lower salivary cortisol and reduced QoL at two years of age. Thus, our study support a role of chronic stress indicated by lower cortisol levels in development of asthma, but with unclear links to atopic dermatitis and allergic sensitisation in young children.

In line with previous studies finding marginally lower cortisol in adolescents with low socioeconomic status (39), we included socioeconomic data as well as second-hand smoking into regression analyses. However, none of these factors were found to be significant confounders, possibly reflecting the overriding effects by atopic diseases in the children, as well as a low frequency of second-hand smoke in our cohort.

### Strengths and limitations

The study strengths include a prospective design of a reasonably large group of children included in infancy with and without acute bronchiolitis and atopic disease, a high rate of follow-up investigations, repeated measurements and stringent clinical characterisation of the subjects. Also, the findings appear robust, as the associations remained significant after relevant adjustments.

The lack of significant associations between QoL and salivary cortisol in the control group may be due to the relatively few subjects with recurrent bronchial obstruction, shown to be most consistently associated with reduced QoL and salivary cortisol, and that the control children may be more heterogeneous, possibly with a lower risk of future asthma development, or that the control children in general had a higher QoL.

As previously reported (7, 8), we decided a priori not to adjust for multiple analyses, as the QoL domains were not independent from each other. Also, the associations with the different QoL domains point in the same direction, limiting the likelihood of incidental findings. The only domain pointing in the opposite direction was Change in health, where a lower score reflects less improvement from an earlier time point. This domain has been shown to be higher in children with chronic diseases than general population children (5), and was the only domain with higher scores in the bronchiolitis than the control group in our study.

The rate of AD and allergic sensitisation was high among the controls, possibly reflecting that parents with some atopic manifestation were more likely to enrol their child into the study.

The use of a single morning salivary cortisol measurement to improve feasibility of a high data collection may be a limitation of our study. However, previous studies of single morning measurements (18) and the lack of significant day-to-day variation between three samples taken at 4-to 8-day intervals (40), suggest that single measures may reflect the habitual morning cortisol state. Also, we sampled as soon as possible after the first awakening after 6:00 a.m. (13), to encompass a possible morning awakening response and the top circadian morning value (25).

## Conclusion

At two years of age QoL was positively associated with morning salivary cortisol in children who had undergone moderate to severe acute bronchiolitis in infancy. The associations were influenced by recurrent bronchial obstruction and, to a limited extent, by atopic dermatitis. The QoL in one-year-old children was associated with salivary cortisol 10 months later.

## Abbreviations

QoL: Health related quality of life
ITQOL: The Infant Toddler Quality of Life Questionnaire
AD: Atopic dermatitis

## CONTRIBUTIONS BY EACH AUTHOR

**Leif Bjarte Rolfsjord, M.D.**, main author, has given substantial contributions to the conception and design of the work, acquisition of the data, analysis and interpretation of the data, drafted the work, finally approved the version to be published and agreed to be accountable for all aspects of the work in ensuring that questions related to the accuracy or integrity of any part of the work are appropriately investigated and resolved.

**Håvard Ove Skjerven, M.D., Ph.D.** is PI of the Bronchiolitis study, has given substantial contributions to the conception and design of the work and contributed to acquisition of the data for the work. He has revised it critically for important intellectual content. He has finally approved the version to be published and agreed to be accountable for all aspects of the work in ensuring that questions related to the accuracy or integrity of any part of the work are appropriately investigated and resolved.

**Egil Bakkeheim, M.D., Ph.D.** has given substantial contributions to the conception and design of the work, especially morning salivary cortisol data and contributed to analysis of the data for the work. He has revised it critically for important intellectual content. He has finally approved the version to be published and agreed to be accountable for all aspects of the work in ensuring that questions related to the accuracy or integrity of any part of the work are appropriately investigated and resolved.

**Teresa Løvold Berents, M.D., Ph.D.** has given substantial contributions to the conception and design of the work and contributed to acquisition of the data for the work, especially of atopic dermatitis data. She has revised it critically for important intellectual content. She has finally approved the version to be published and agreed to be accountable for all aspects of the work in ensuring that questions related to the accuracy or integrity of any part of the work are appropriately investigated and resolved.

**Kai-Håkon Carlsen, M.D., Ph.D.** has given substantial contributions to the analysis and interpretation of the data for the work. He has revised it critically for important intellectual content. He has finally approved the version to be published and agreed to be accountable for all aspects of the work in ensuring that questions related to the accuracy or integrity of any part of the work are appropriately investigated and resolved.

**Karin C. Lødrup Carlsen, M.D., Ph.D.** has given substantial contributions to the conception, design, analysis and interpretation of the data of the work. She has given substantial contributions to drafting the work and has revised it critically for important intellectual content. She has finally approved the version to be published and agreed to be accountable for all aspects of the work in ensuring that questions related to the accuracy or integrity of any part of the work are appropriately investigated and resolved.

## ACKNOWLEDGEMENTS

We warmly acknowledge all participating children and parents and the members of the Bronchiolitis Study Group, and study nurses, see Supporting Information, and the several hundred study staff that were involved in recruiting patients and running the study. Warm thanks also to Johan Alm and Ann-Christine Sjöbeck, Department of Clinical Science and Education, Karolinska Institutet, Stockholm, Sweden, for the analysis of the salivary cortisol samples.

